# Differences in Activation of Intracellular Signalling in Primary Human Retinal Endothelial Cells Between Isoforms of VEGFA-165

**DOI:** 10.1101/2019.12.12.871947

**Authors:** Wendelin Dailey, Roberto Shunemann, Fang Yang, Megan Moore, Austen Knapp, Peter Chen, Mrinalini Deshpande, Brandon Metcalf, Quentin Tompkins, Alvaro E. Guzman, Jennifer Felisky, Kenneth P. Mitton

## Abstract

**Purpose:** There are reports that a b-isoform of Vascular Endothelial Growth Factor-A-165 (VEGFA_165_b) is predominant in normal human vitreous, switching to the a-isoform (VEGFA_165_a) in the vitreous of some diseased eyes. While these isoforms appear to have a different ability to activate the VEGF-Receptor-2 (VEGFR2) in various endothelial cells, the nature of their ability to activate intracellular signalling pathways is not fully characterized, especially in retinal endothelial cells. We determined their activation potential for two key intracellular signalling pathways (MAPK, AKT) over complete dose-response curves and compared potential effects on the expression of several VEGFA_165_ target genes in primary human retinal microvascular endothelial cells (HRMECs).

**Methods:** To determine full dose-response curves for the activation of MAPK (ERK1/2), AKT and VEGFR2, direct in-cell western assays were developed using primary Human Retinal Microvascular Endothelial Cells (HRMECs). Potential differences in dose-response effects on gene expression markers related to endothelial cell / leukocyte adhesion (*ICAM1, VCAM1* and *SELE*) and tight-junctions (*CLDN5* and *OCLN*) were tested by quantitative-PCR.

**Results:** Activation dose-response analysis revealed much stronger activation of MAPK, AKT and VEGFR2 by the a-isoform at lower doses. MAPK activation in primary HRMECs displayed a sigmoidal dose-response to a range of VEGFA_165_a concentrations spanning 10-250 pM, which shifted higher into the 100-5,000 pM range with VEGFA_165_b. Similar maximum activation of MAPK was achieved by both isoforms at high concentration. Maximum activation of AKT by VEGFA_165_b was only half of the maximum activation from VEGFA_165_a. At a lower intermediate dose, where VEGFA_165_a activated intracellular signalling stronger than VEGFA_165_b, the changes to VEGFA target gene expression was generally greater with VEGFA_165_a.

**Conclusions:** In primary HRMECs, VEGFA_165_a could maximally activate MAPK and AKT at lower concentrations where VEGFA_165_b had relatively little effect. The timing for maximal activation of MAPK was similar for both isoforms, which is different than reprorted for non-retinal endothelial cells. While VEGFA_165_a and VEGFA_165_b are limited to the sequence of their six C-terminal six amino acids, this results in a large difference in their ablility to activate at least two key intracellular signalling pathways and potentially VEGF target gene expression in primary human retinal endothelial cells.

## Introduction

Vascular Endothelial Growth Factor-A-165 (VEGFA_165_) is the isoform of VEGF that is primarily responsible for driving retinal vascular pathology in diabetic retinopathy (DR), age-related macular degeneration (AMD) and retinopathy of prematurity (ROP), through activation of VEGF Receptor-2 (VEGFR2). A seminal analysis of the vitreous fluids of patients with diabetic retinopathy established the presence of elevated VEGF concentrations associated with this and other conditions [1]. That study contributed to the eventual development of intravitreal VEGF-blocking drugs to treat neovascularization and edema in AMD, DR, and the current exploration of their use for ROP [2–5].

Blockade of VEGFA activity is provided by the use of intravitreal drugs including ranibizumab (Lucentis, Genentech), bevacizumab (Avastin, Genentech), pegaptanib (Macugen, Bausch & Lomb), and aflibercept (Eylea, Regeneron). Some VEGF-blocking drugs injected into the vitreous can enter the systemic circulation and, in the case of bevicizumab and aflibercept they can substantially decrease serum VEGF concentration for several days [6]. There is interest to advance VEGF-regulating therapies by more precise titration of VEGF concentration or modulating specific VEGF-mediated signalling rather than the full blockade provided by current drugs. This field will continue to benefit from a more complete knowledge of VEGF’s signaling mechanisms within retinal endothelial cells. Unfortunately, much previous VEGF research does not technically differentiate between VEGFA’s isoforms and in the case of the retina there is far less investigation using retinal endothelial cells, and even less from human retina.

In addition to endothelial cells, VEGF receptors are expressed by cells of the immune system, including early and late hematopoietic progenitor cells, dendritic cells, T-lymphocytes and macrophages [7]. Among the three VEGF receptor tyrosine-kinases (R1, R2, R3), VEGFR2 is required for angiogenesis and vasculogenesis, and its expression is most abundant in endothelial and endothlial progenitor cells [8–10]. VEGFR2 binds several isoforms of the VEGFA that are produced from alternative-splicing [11–14]. The most frequently detected isoforms are VEGFA_121_, VEGFA_165_ and VEGFA_189_, with 121, 165 and 189 amino acids respectively. VEGFA_121_ is most diffusible, lacking the heparin-binding domains of VEGFA_165_ and VEGFA_189_. VEGFA_189_ has even more affinity for heparin than VEGFA_165_ with an additional heparin-binding domain encoded by exon-6 [11,12]. VEGFA_165_ bind as a dimer to activate VEGFR2, and it also binds Neuropilin-1 as a co-receptor. VEGFA_121_ lacks two regions required to binding Neuropilin-1.

A vast knowledge of VEGF-mediated signalling is mostly derived from the study of non-retinal cell types, and is beyond the scope of this paper but readers can refer to the excellent review by Koch and Claesson-Welch [15]. In endothelial cells, several pathways are activated by VEGFA_165_ that affect proliferation, migration, survival and endothelial permeability. Two of these pathways include MAPK (ERK1/2) and the AKT (PKB), where MAPK is a dominant regulator of cell proliferation and AKT modulates cell survival and permeability [16–19]. Autophosphorylation of VEGFR2 (Y1175) leads to RAS-independent activation of the PLCγ (Phospholipase-Cγ) / Protein Kinase C / MAPK pathway [20,21]. Activation of VEGFR2 also leads to activation of the TSAD / PI3K (Phosphotidylinositol-3 Kinase) / PDK (Phosphoinositide-dependent protein kinase) / AKT pathway [22,23]. AKT phosphorylates the BCL-2-associated death protein (BAB) and Caspase-9 to block apoptosis and increase EC-survival [24]. AKT also activates eNOS (endothelial Nitric-Oxide Synthase, *NOS3)* to modulate vascular permeability [25].

More recently b-isoforms of VEGFA were described that differ from the previously known a-isoforms in the sequence of their C-terminal six amino acids: CDKRPP in VEGFA_165_a becomes SLTRDKD in VEGFA_165_b [13,14]. This difference renders VEGFA_165_b incapable of bringing Neuropilin-1 into a receptor/ligand complex with VEGFR2 [26–28]. Similar to VEGFA_121_, VEGFA_165_b is reported to be less angiogenic than VEGFA_165_a [13,29,30]. VEGFA_165_b’s binding affinity for VEGFR2 itself is similar to that of VEGFA_165_a in Human Umbilical Vein Endothelial Cells (HUVECs) and less than VEGFA_165_a in direct binding anlaysis in transformed HEK293 cell assays [31,32]. VEGFA_165_b also displays weaker activation of VEGFR2 and ERK1/2 (MAPK) than VEGFA_165_a in transfected porcine aortic endothelial (PAE) cells [31]. Perrin et al. (2005), reported a switch from mostly VEGFA_165_b to mostly VEGFA_165_a in the vitreous of patients with active diabetic retinopathy [33]. A similar shift, from VEGFA_165_b to VEGFA_165_a, was reported in vitreous humor from young patients with ROP [34]. A murine equivalent of isoform switching (VEGFA_164_b to VEGFA_164_a) was also reported in a mouse model of oxygen-induced retinopathy [35]. Several publications using isoform specific antibodies have detected VEGFA_165_b in human and mouse tissues, but their presense from RNA analysis in sheep hypothalamus is controversial [36].

While single-dose studies using transfected cells or non-retinal endothelial cells indicate that VEGFA_165_b activates VEGFR2 and MAPK less than VEGFA_165_a, we do not know if this is true throughout a full dose range. We also wanted to determine full-dose response curves for activation of intracellular signaling in primary human retinal microvascular endothelial cells. We hypothesized that there is a significant difference in the full dose-reponse curves for activation of these pathways in human retinal endothelial cells between VEGFA_165_a and VEGFA_165_b. To address this, we used *in situ* assays to determined full dose-reponse curves for the activation of MAPK (ERK1/2), AKT and VEGFR2 in primary HRMECs for VEGFA_165_a and VEGFA_165_b. Our results demonstrate that there is a significant dose-response difference in the ability of these two isoforms to activate intracellular signalling kinases and that these dose-response differences propagate to differences in VEGF-target gene expression in primary human retinal endothelial cells.

## Methods

### Certification of Primary Human Retinal Microvascular Endothelial Cells

Primary Human Retinal Microvascular Endothelial Cells (HRMECs, genotype XY) were obtained from Cell Systems as passage-3 cells (Kirkland, WA, Catalog number: ACBRI-181). Cells used for experimentation were not used past passage-7. Endothelial character of the cell line was established by Cell Systems immunofluorescence testing: > 95% positive by immunofluorescence for Cytoplasmic VWF / Factor VIII, Cytoplasmic uptake of Di-I-Ac-LDL, Cytoplasmic CD31, and < 1% by immunofluorescence for GFAP (Glial Fibrillary Acidic Protein), Glutamine Synthetase, NG2 (Neural/Glial Antigen 2), and PDGFR-beta (Platelet Derived Growth Factor Receptor beta). The cell line was also subjected to short tandem repeat (STR) Profile Testing by Cell Systems at the Master Level (P1), performed by the laboratory analysis service of the American Type Culture Collection (ATCC). Seventeen STR-loci plus the gender determining locus, Amelogenin, were amplified using the commercially available PowerPlex® 18D Kit from Promega. The cell line sample was processed using the ABI Prism® 3500xl Genetic Analyzer. Data were analyzed using the GeneMapper® IDX v1.2 software (Applied Biosystems). Appropriate positive and negative controls were used throughout the test procedure. DNA Profile (STR #12625 Analysis) was as follows: TH01: 9, 9.3; D5S818: 9, 13; D13S317: 8, 10; D7S820: 11, 12; D16S539: 11, 12; CSF1PO: 10, 12; Amelogenin: X, Y; vWA: 16, 19; TPOX: 8, 11. The ATCC test conclusions were that the submitted sample profile is human, but not a match any profile in the ATCC STR database, as would be expected for primary HRMECs. Additionally, routine testing for retinal endothelial character in the author’s laboratory (Eye Research Institute, Oakland University) confirmed that the HRMECs express all three of the human Norrin receptor-complex genes *FZD4, LRP5* and *TSPAN12*, using Taqman real-time PCR analysis (data not shown). HRMECs also demonstrated VEGF-medidated regulation of several VEGF target genes, as shown in the results.

### Cell Culture and Antibodies

EndoGRO-MV Complete media kit was obtained from Millipore (Burlington, MA). EndoGro basal medium was supplemented with rhEGF, L-Glutamine, Heparin Sulfate and Ascorbic Acid according to the kit instructions. Additionally, supplementation with FBS and Hydrocortisone Hemisuccinate (1μg/ml) was dependent upon the assay requirements. Recombinant human VEGFA_165_a and VEGFA_165_b were obtained from R&D Systems (Minneapolis, MN). Odyssey Blocking Buffer and an Odyssey Infrared Imager were purchased from LI-COR Biosciences (Lincoln, NE). The antibodies used for in situ labeling of HMRECs (In Cell Westerns, ICW) and Western blotting are summarized in Table 1. Mini Protean TGX gels were purchased from Bio-Rad (Hercules, CA).

**Table-1.** Specific Antibodies used and Sources

| **Antibody** | **IgG Species** | **Company** | **Cat. No.** |
| --- | --- | --- | --- |
| Phospho-AKT (P-Ser473) | Rabbit | Cell Signaling Technologies | 4058 |
| AKT(pan) | Rabbit | Cell Signaling Technologies | 4691 |
| Phospho-p44/42 MAPK (Thr202/204) | Rabbit | Cell Signaling Technologies | 4370 |
| p44/42 MAPK (pan) | Rabbit | Cell Signaling Technologies | 4695 |
| Phospho-VEGFR2 (Tyr1175) | Rabbit | Cell Signaling Technologies | 2478 |
| Beta-Actin | Mouse | Millipore | MAB1501 |
| Anti-mouse IgG IRDye 800CW | Goat | LI-COR | 32210 |
| Anti-rabbit IgG IRDye 680RD | Goat | LI-COR | 68071 |
| Anti-rabbit IgG IRDye 800CW | Goat | LI-COR | 32211 |
| Anti-mouse IgG IRDye 680RD | Goat | LI-COR | 68070 |

### Immunoblotting of activated MAPK and AKT in HRMECs

HRMECs were grown in 100mm dishes that had been pre-coated with Attachment Factor (Cell Systems, WA). Media was EndoGRO (No VEGF, Millipore Sigma, Burlington, MA) with 5% FBS to establish confluency. When cells were confluent, the media was replaced with fresh media with or without VEGFA_165_ isoforms. After incubation for 10 minutes, the cells were washed with ice cold PBS and the dishes were scraped to detach the cells. The cell suspension was centrifuged at 1000 X g for 5 minutes and the cells were reconstituted in RIPA cell-lysis buffer (150 mM NaCl, 1% Triton-X-100, 0.1% SDS, 50 mM Tris, pH 8.0, 10 mM Sodium Fluoride, 1 mM Sodium Orthovanadate and complete Protease inhibitor cocktail (1 tablet/10 ml)). The cells were sonicated or vortexed every 10 minutes while kept on ice for 30 minutes. The cell lysate was collected after centrifuging at 14,000 X g for 15 minutes at 4°C. The protein concentration was measured using Pierce BCA Protein Assay (Thermo Scientific). The samples were prepared with Laemmli sample buffer and loaded onto a (4-15%) gradient gels for SDS-PAGE electrophoresis. After electrophoresis the gels were equilibrated in cold transfer buffer and transferred to PDVF membranes overnight. The membranes were blocked with Odyssey blocking buffer and incubated with appropriates rabbit primary antibodies (Table-1) along with mouse monoclonal Actin antibody which was used for normalization. After washing with PBS, 0.1% tween-20 (4 × 5 min), the membranes were incubated with secondary antibodies (Goat Anti-rabbit IRDye680 and Goat Anti-mouse IRDye800CW) for 30-60 minutes. Membranes were washed and scanned on an Odyssey Infrared imager (Li-Cor, Lincoln, NE).

### Dose-response analysis of intracellular signaling in primary HMRECs

HRMECs were seeded into black 96 well plates (5000/well) that had been coated with Attachment Factor. The cells were grown to confluence using fully supplemented EndoGRO-MV media for 4-5 days. The cells were serum starved overnight using EndoGRO-MV without Hydrocortisone. The cells were treated with VEGFA_165_a or VEGFA_165_b for varied lengths of time after which the treatments were immediately removed and replaced with of 4% Paraformaldehyde. The cells were fixed for 20 minutes at room temperature followed by permeabilization with PBS, 0.1% Triton X-100 (10 minutes). Cells were blocked by incubation with Odyssey Blocking Buffer (Li-Cor) for 1.5 hours at room temperature and incubated with primary antibodies (1:200) for either 2 hours at room temperature or over night at 4 °C. Rabbit antibodies were used against the proteins of interest and a mouse monoclonal anti-Beta-Actin antibody was used for normalization. The cells were washed with PBS, 0.1% Tween 20 (5×5 minutes) and incubated with secondary antibodies, Goat anti-Rabbit IRDye 800CW & Goat anti-Mouse IRDye 680RD (1:750) for 45 minutes at room temperature. After washing with PBS 0.1% Tween 20 (5×5 minutes) the plates were scanned on an Odyssey Imager (Li-Cor). For time-response and dose-response experiments, the doses of VEGFA_165_a and VEGFA_165_b were assayed in quadruplicate wells and dose-response experiments were repeated three times to confirm reproducibility of relative dose-response curves for activation of MAPK (phospho-Thr202/Tyr204), AKT (phospho-Ser473) and VEGFR2 (phospho-Tyr1175). Data were fit to a 4-parameter log-logistic response function Eq (1) at the 95% confidence level for each dose *x* using the Drc package for R. [37]. The parameters fit were: *b*, the steepness of the curve at *e* the effective dose ED_50_, with *c* and *d* the lower and upper limits of the response. The 95% confidence level for fitting was used to produce curves.

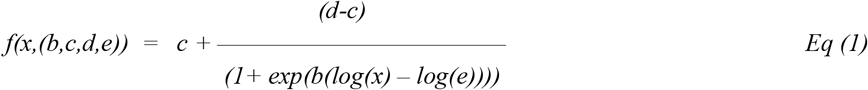

### Quantitative PCR analysis of HMREC Gene Expression

Primary HMRECs were grown to confluence in six well plates. After the desired treatment, cells were trypsinized. Total RNA was isolated using the Absolutely RNA miniprep kit (Agilent, Santa Clara, CA). Cells were homogenized in 200μL of lysis buffer. Homogenization was accomplished using conical pellet-pestles in 1.5 mL microfuge tubes, with a hand-held rotary tool (Bel-Art,Wayne, NJ). First-strand cDNA was synthesized by reverse transcribing 500 ng of total RNA per sample using either the AffinityScript qPCR DNA Synthesis kit (Agilent), or the LunaScript RT Super Mix Kit (NEB # E3010(S/L), Ipswich, MA) with Oligo-dT priming. The reaction conditions were according to the manufacturer’s instructions. For AffinityScript: 25°C for 5 min, 42°C for 20 min, 95°C for 5 min, and 10°C for 10 min. For LunaScript RT: 25°C for 2 min, 55°C for 10 min, and 95°C for 1 min. All compared samples were processed using the same reagent set. Stock first strand cDNA preparations were stored at −70°C, and were not used for analysis after a maximum of three freeze-thaws. Duplex reaction format was utilized with FAM-labeled probe/primer pairs for the gene of interest and VIC-labeled probe/primer-limited pairs for TBP (Tata-Binding Protein) as the normalizer gene (ThermoFisher, Waltham, MA). For real-time PCR reactions, sample first-strand cDNA was diluted 5-fold with deionized water and 2μL added to 18μL of master mix for 20μL PCR reactions. Triplicate reactions were used for each sample. Master Mix chemistries were either 2x Gene Expression Master Mix (ThermoFisher, Applied Biosystems # 4369016, Waltham, MA), or the Luna Universal Probe qPCR Master Mix (2x) Gene Expression (New England BioLabs # M3004L, Ipswich, MA), both mixes with the Rox reference dye option. Reactions were run on either an Mx3000P real-time PCR system using the MxPro software, or an AriaMx Real-time PCR System using the AriaMx HRM QPCR Software (Agilent, *Santa Clara, CA*). Gene expression assays were evaluated for high PCR efficiency using a dilution series of HMREC cDNA to ensure validity of using the delta-delta Ct method for comparing relative gene expression. Each replicate reaction was internally normalized relative to endogenous TBP gene expression. The specific assay probe sets used for gene expression analysis are listed in Table 2

**Table 2.** Taqman gene expression probes for HRMEC gene expression.

| Gene | Probe Set # | Spans Exons |
| --- | --- | --- |
| *CLDN5* | Hs01561351_m1 | 1-2 |
| *ICAM1* | Hs00164932_m1 | 2-3 |
| *OCLN* | Hs00170162_m1 | 5-6 |
| *SELE* | Hs00174057_m1 | 4-5 |
| *TBP* | Hs00427620_m1 | 3-4 |
| *VCAM1* | Hs01003372_m1 | 6-7 |

## ESULTS

### Preliminary observations

The cell-based experimental results reported here were inspired by observations made during preliminary testing for a different project to develope an intravitreal injection model using VEGFA_165_a to cause dilation of primary retinal veins, which was reported to occur in the Long Evans rat [38]. That model typically involves a high dose of VEGFA. Using fluoresceine angiography, and SD-OCT imaging, it was noted that similar effects on the vasculature could be induced with VEGFA_165_b or VEGFA_165_a. While the observational studies did not use large numbers of rats, they are included as supplimental observations because they brought our attention that the dose-responses for intracellular pathway activation were not fully known in primary human retinal endothelial cells.

### Activation of MAPK and AKT in primary HRMECs by VEGFA_165_a and VEGFA_165_b

To examine the effect of VEGF_165_ isoforms on intracellular signaling pathways we chose primary human retinal microvascular endothelial cells (HRMECs) and initially used immunoblotting to test activation of MAPK and AKT pathways by both isoforms. These pathways were examined as two of the intracellular signaling kinases implicated in mediating the effects of VEGFA on endothelial proliferation and blood-retinal barrier. Using a high dose (100 ng/mL, 5300 pM) for maximal activation, both isoforms of VEGFA_165_ increased the amounts of the active form of MAPK (Figure-1a) and the active form of AKT (Figure-1b). From three separate full experimental repeats, the average activation was about 2.7-fold and 1.6-fold (Figure-1c) for MAPK and AKT respectively. While VEGFA_165_a and VEGFA_165_b treatments always increased the activation of both kinases, the magnitude of the increase was variable between experiments as illustrated by comparing two of the three experiments on the same gel (Figure 1a,b).

**Figure 1.**
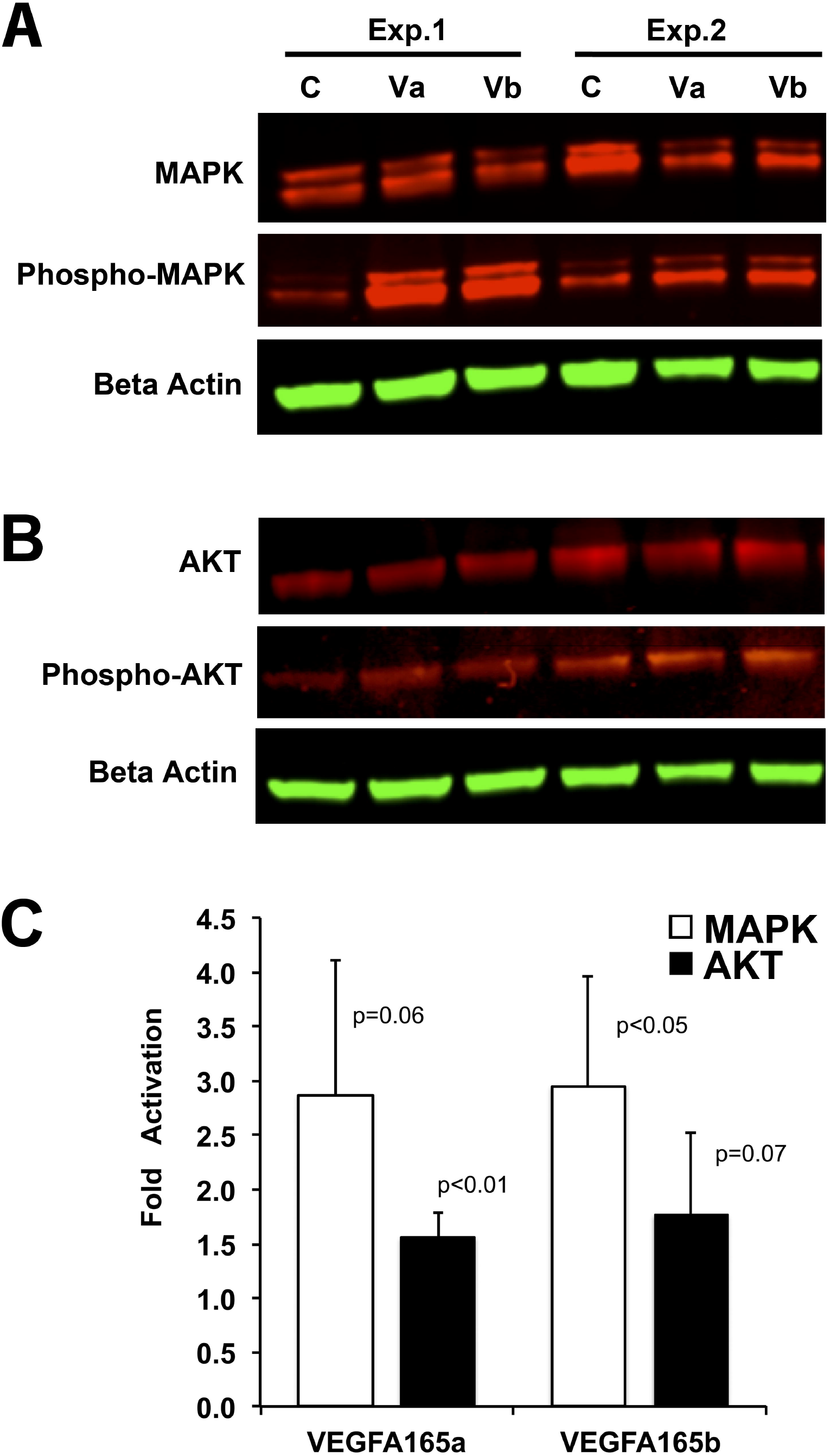
MAPK and AKT are activated in HRMECs by VEGFA_165_a and VEGFA_165_b. **A)** Immunoblots of total MAPK and activated MAPK (Phospho-MAPK) 10 minutes after treatment with VEGFA_165_a (Va), VEGFA_165_b (Vb), or media control (C). Two representative experiments are shown. **B)** Immunoblots of total AKT and activated AKT (Phospho-AKT) 10 minutes after treatment with VEGFA_165_a (Va), VEGFA_165_b (Vb), or media control (C). Two representative experiments are shown. **C)** Fold activation of MAPK and AKT from three immunoblotting experiments. P-values are shown, t-test relative to media controls (1-fold).

### Dose-Response for the Activation of MAPK by VEGFA_165_a and VEGFA_165_b

While repeated immunoblotting experiments confirmed activation of MAPK and AKT by both VEGFA_165_ isoforms, the experimental variability introduced by protein extraction, protein assays, electro-transfer, band-shape and image analysis was judged to be unsuitable for dose-response analysis. To address the variability, new *in situ* assays, or In-Cell Western (ICW) assays, were developed using the same kinase-specific antibodies while removing the sample processing required for immunoblotting. This also permitted faster fixation of cells at the desired time, the use of many doses, and the use of several biological replicates per dose.

Activation of the MAPK (ERK1/2) pathway in primary HRMECs occurred maximally at 10 minutes for both VEGFA_165_a and VEGFA_165_b treatment, and returned to control levels by 90 minutes, using a dose of 2ng/mL (100 pM). See Figure-2a. The activation of VEFGA_165_a was significantly stronger than that of VEGFA_165_b at 10 minutes (t-test, P<0.01).

**Figure 2.**
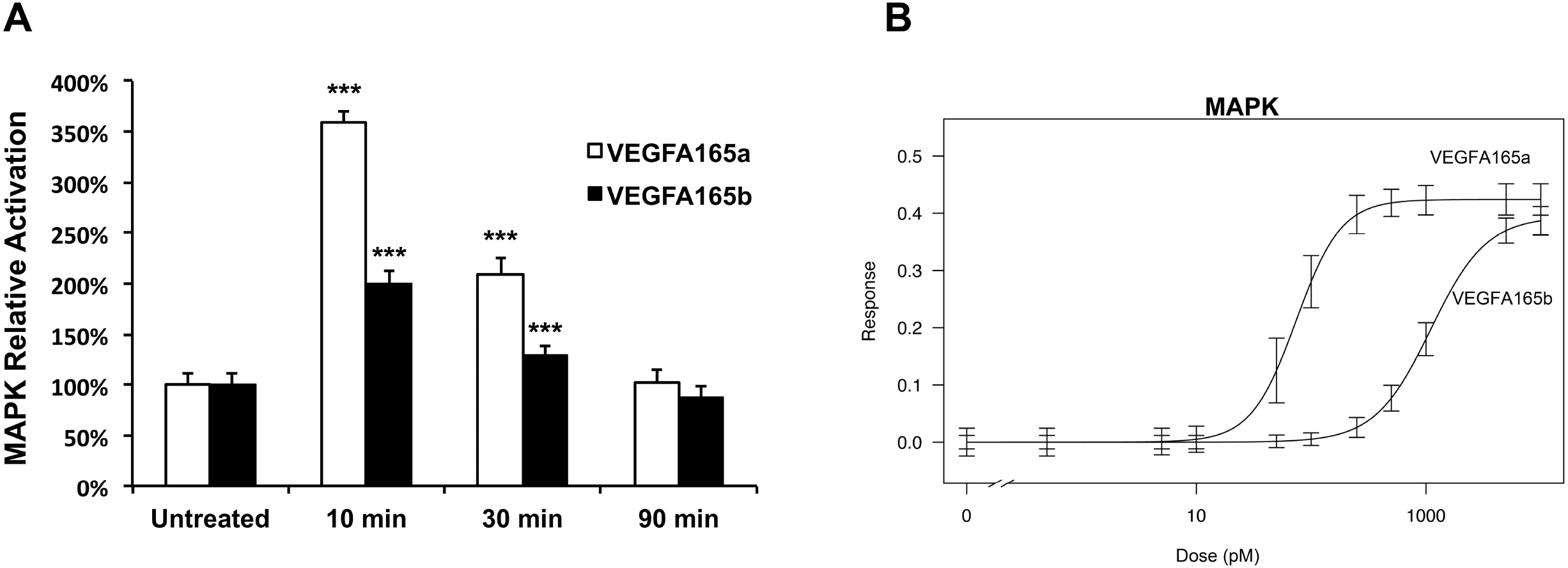
VEGFA_165_a strongly activates MAPK in primary HRMECs at doses where VEGFA165b has little effect. Activation of MAPK was measured by *in-situ* immunofluorescence with a phospho-specific antibody (phospho-Thr202/Tyr204) for p44/42. **A)** Activation of MAPK at three time-points after the addition of 100 pM VEGFA_165_a or VEGFA_165_b. Bar shows standard deviation. (t-test, *** p<0.001 relative to untreated control, n=8 biological replicates.) **B)** Dose-response curves for activation of MAPK. The ED_50_ was 73 pM for VEGFA_165_a and 1015 pM for VEGFA_165_b. Bars indicate the 95% confidence interval for data fit to the 4-parameter log-logistic function. N=4 biological replicates per dose.

We next carried out dose-response analysis for the activation of the MAPK at 10 minutes with cells pre-adapated to lower serum. Eleven doses of each isoform, with 4 biological replicates per dose, were evaluated and dose-response data for MAPK activation were fit to the Log-Logistic 4-parameter dose-response function using the Drc package in R [37]. (Figure-2b) Activation of MAPK by VEGFA_165_b showed a typical sigmoidal dose-response. Compared to VEGFA_165_b, the dose-response curve for VEGFA_165_a displayed a strong allosteric shift to a more binary-like activation response. VEGFA_165_a was more potent for activation of MAPK at lower doses. A similar maximum level of MAPK activation could be achieved with the highest dose tested (10,000 pM) using either isoform. Activation of MAPK by VEGFA_165_b was only 10% of that generated by VEGFA_165_a using a dose of 250 pM. The ED_50_ was 73 pM for VEGFA_165_a and 1015 pM for VEGFA_165_b for the experiment shown. The experiment was repeated three times confirming the large difference in ED_50_ between the two isoforms. Differences between the ED_50_ values were typically in the 800 to 1000 pM range.

### Dose-Response for the Activation of AKT Activation by VEGFA_165_a and VEGFA_165_b

Maximum activation of the pAKT pathway in HRMECs occurred 30 minutes after VEGFA_165_a treatment and 15 minutes after VEGFA_165_b treatment (Figure-3a) using what was thought to be a maximally activating dose (100 ng/mL 5,300 pM). Dose response analysis was carried out for AKT activation using these respective maximum time points and the data were fit to the 4-parameter log-logistic response function. (Figure-3b). With VEGFA_165_a, AKT activation in HRMECs displayed a much steeper dose-response curve than for VEGFA_165_b. The ED_50_ values were 53 pM for VEGFA_165_a compared to 126 pM for VEGFA_165_b. The maximum activation of AKT generated by the b-isoform was less than half that obtained from treatment with the a-isoform. Repetition of the experiment confirmed similar relative responses.

**Figure 3.**
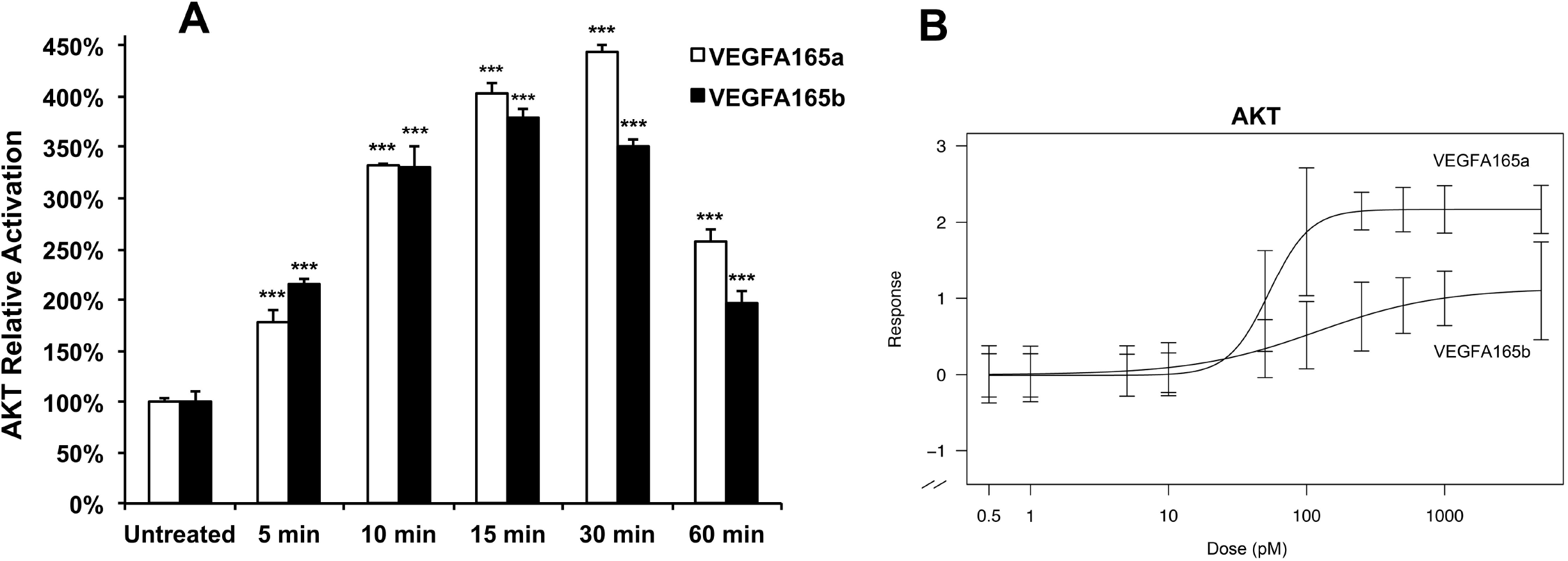
VEGFA_165_b is a weaker activator of AKT in primary HRMECs compared to VEGFA165a. **A)** Activation of AKT was measured by *in-situ* immunofluorescence with a phospho-specific antibody for Ser473-AKT at five time-points after the addition of 20 ng/mL (1,250 pM) VEGFA_165_a or VEGFA_165_b. Bar shows standard deviation. (t-test, ***p<0.001 relative to untreated control, n=4 biological replicates.) **B)** Dose response curves for activation of AKT. The ED_50_ value for AKT activation by VEGFA_165_a was 53 pM compared to 126 pM for VEGFA_165_b. Bars indicate the 95% confidence interval for data fit to the 4-parameter log-logistic function. N=4 biological replicates per dose.

### Dose-Response for the Activation of VEGFR2 by VEGFA_165_a and VEGFA_165_b

To test for relative difference in dose-response at the level of the receptor itself we examined the dose-response for activation of VEGFR2. Maximum activation of the receptor was fast, within less than a few minutes, as quickly as cells could be processed for treatment and fixation. Five and ten minute experiments were similar in relative response, so ten minutes were used for the dose-response experiments and data were fit to the 4-parameter log-logistic response function as shown in Figure-4a. VEGFA_165_a treatment achieved maximum activation by about 500 pM whereas VEGFA_165_b caused little if any activation at that dose. The ED_50_ for activation by VEGFA_165_a was 254 pM compared to 1192 pM for the VEGFA_165_b. Furthermore, the maximum activation by VEGFA_165_b was 73% of that obtained with VEGFA_165_a. Again, the relative pattern was confirmed by a repeated experiment (data not shown).

**Figure 4.**
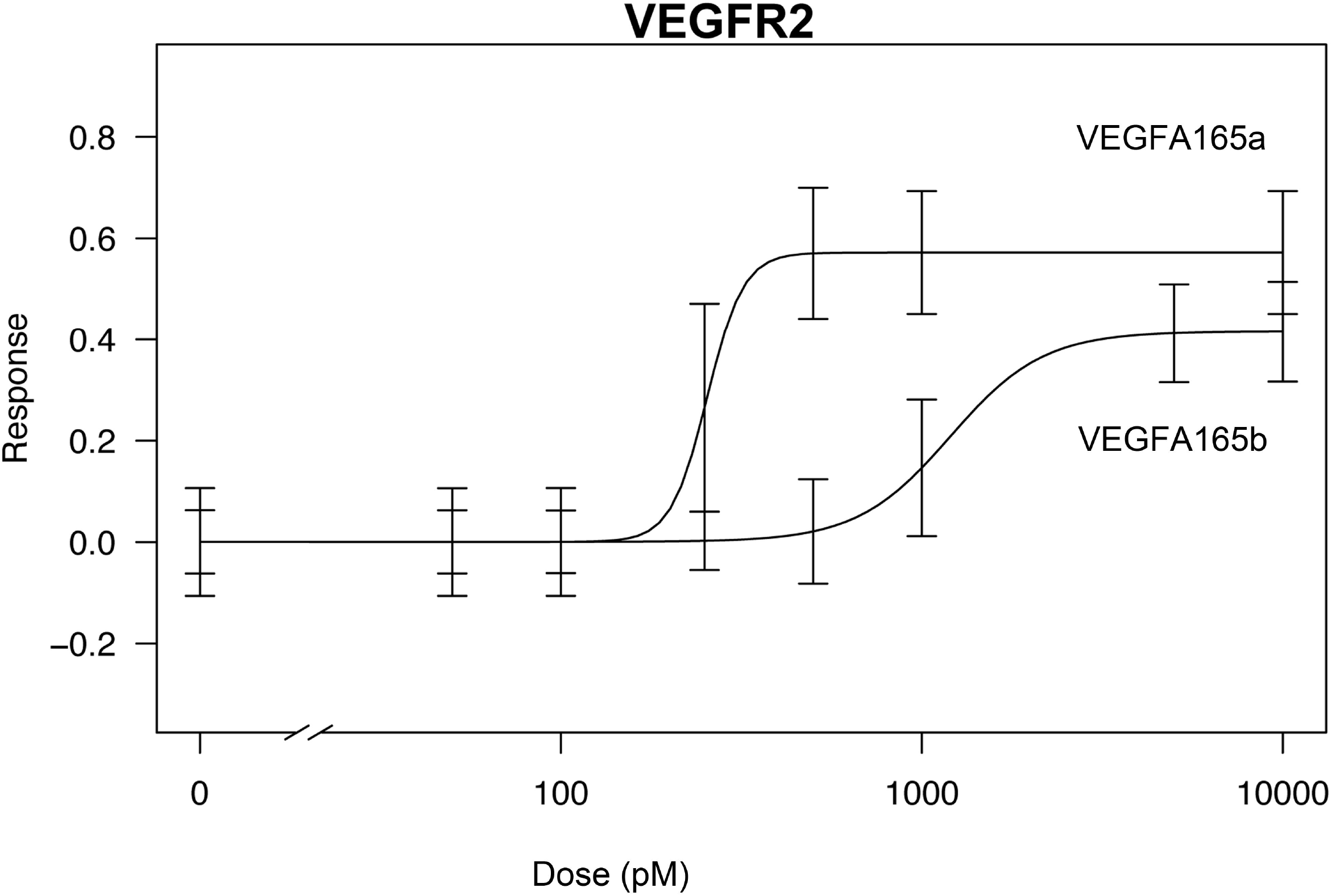
Differences in the activation of intracellular signalling between VEGFA_165_a and VEGFA165b in primary HRMECs begin at the receptor VEGFR2. VEGFA_165_a was a stronger activator of VEGFR2 compared to VEGFA_165_b. The ED_50_ for activation by VEGFA_165_a was 254 pM compared to 1192 pM for VEGFA_165_b. Bars indicate the 95% confidence interval for data fit to the 4-parameter log-logistic function. N=4 biological replicates per dose.

### Isoform Effects on Expression Leukocyte Docking Protein Genes

We first examined gene expression from 1 to 24 hours after treatment with high doses (100 ng/mL, 5300 pM) of VEGFA_165_a and VEGFA_165_b to determine an optimal time-point for monitoring any effects on the expression of *ICAM1, Selectin-E*, and *VCAM1*. A dose of 5300 pM was likely a saturating concentration and more than what would be experienced *in vivo*, to find a suitable time-point where gene expression could be subsequently compared using physiologically relevant doses. All three of these genes displayed increased expression after treatment with both isoforms of VEGFA_165_ (Figure-5 a, c, e). These trends were confirmed by repeated experiments (data not shown). *Of the genes examined here, ICAM1’s* increasing expression trend was more variable in timing, not always maximum at 6 hours, sometimes later by 24 hours (data not shown).

**Figure 5.**
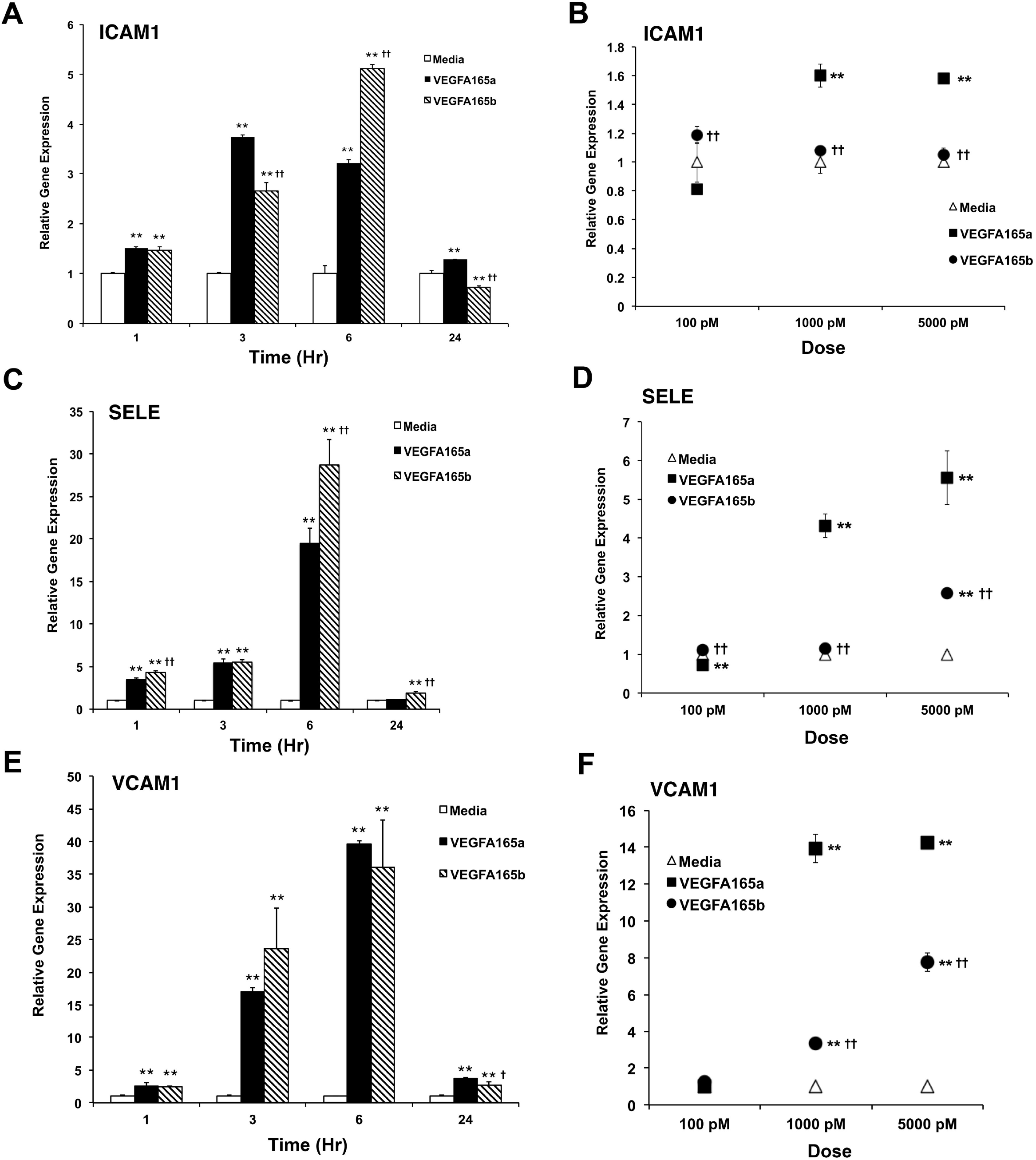
VEGFA165a has a stronger effect on leukocyte-endothelial adhesion gene expression in primary HRMECs than VEGFA165b. Relative gene expression measured by qPCR. **A)** *ICAM1* gene expression at 1, 3, 6 and 24 hours after VEGFA_165_ treatment. Confluent HRMECs were treated with a saturating high-dose of VEGFA_165_a or VEGFA_165_b (100 ng/mL, 5300 pM). **B)** *ICAM1* gene expression comparing VEGFA_165_a and VEGFA_165_b at a low, intermediate and high dose measured after 6-hours. **C)** *Selectin-E* gene expression at 1, 3, 6 and 24 hours after VEGFA_165_ treatment. **D)** *Selectin-E* gene expression comparing VEGFA165a and VEGFA165b at a low, intermediate and high doses measured after 6-hours. **E)** *VCAM1* gene expression at 1, 3, 6 and 24 hours after VEGFA_165_ treatment. **F)** *VCAM1* gene expression comparing VEGFA_165_a and VEGFA_165_b at a low, intermediate and high dose measured after 6-hours. (Triplicate assays, error bars show standard deviation. Anova: compared to media control *p<0.05, **p<0.01; compared to VEGFA_165_a ^†^ p<0.05, ^††^p<0.01)

Selecting a fixed treatment time of 6-hours, and guided by the activation dose-response curves for MAPK and AKT, we tested a low (2ng/mL, 100 pM), intermediate (19 ng/mL, 1000 pM) and high (95 ng/mL, 5000 pM) dose to test for differences in the effects on leukocyte-docking gene expression. For all three leukocyte docking protein genes (*ICAM1, SELE, VCAM1*), VEGFA_165_a increased their expression at the intermediate dose of 1000 pM where VEGFA_165_b still had very little effect (Figure-5 b, d, f).

### Isoform Effects on Expression of Tight Junction Protein Genes

The effects of VEGFA_165_a and VEGFA_165_b on the expression of the two key tight-junction protein genes, CLDNS and OCLN, were also compared. These genes encoding the tight-junction proteins Claudin-5 and Occludin. (See Figure-6) Both isoforms of VEGFA_165_ reduced the expression of these genes by the 6 hour time-point (Figure-6 a, c). We next comparing a low (2ng/mL, 100 pM), intermediate (19 ng/mL, 1000 pM) and high saturating dose (95 ng/mL, 5000 pM) at the fixed 6-hour time point. VEGFA_165_b was less effective at suppressing the expression of *Claudin-5* and *Occludin* compared to VEGFA_165_a (Figure-6 b, d)

**Figure 6.**
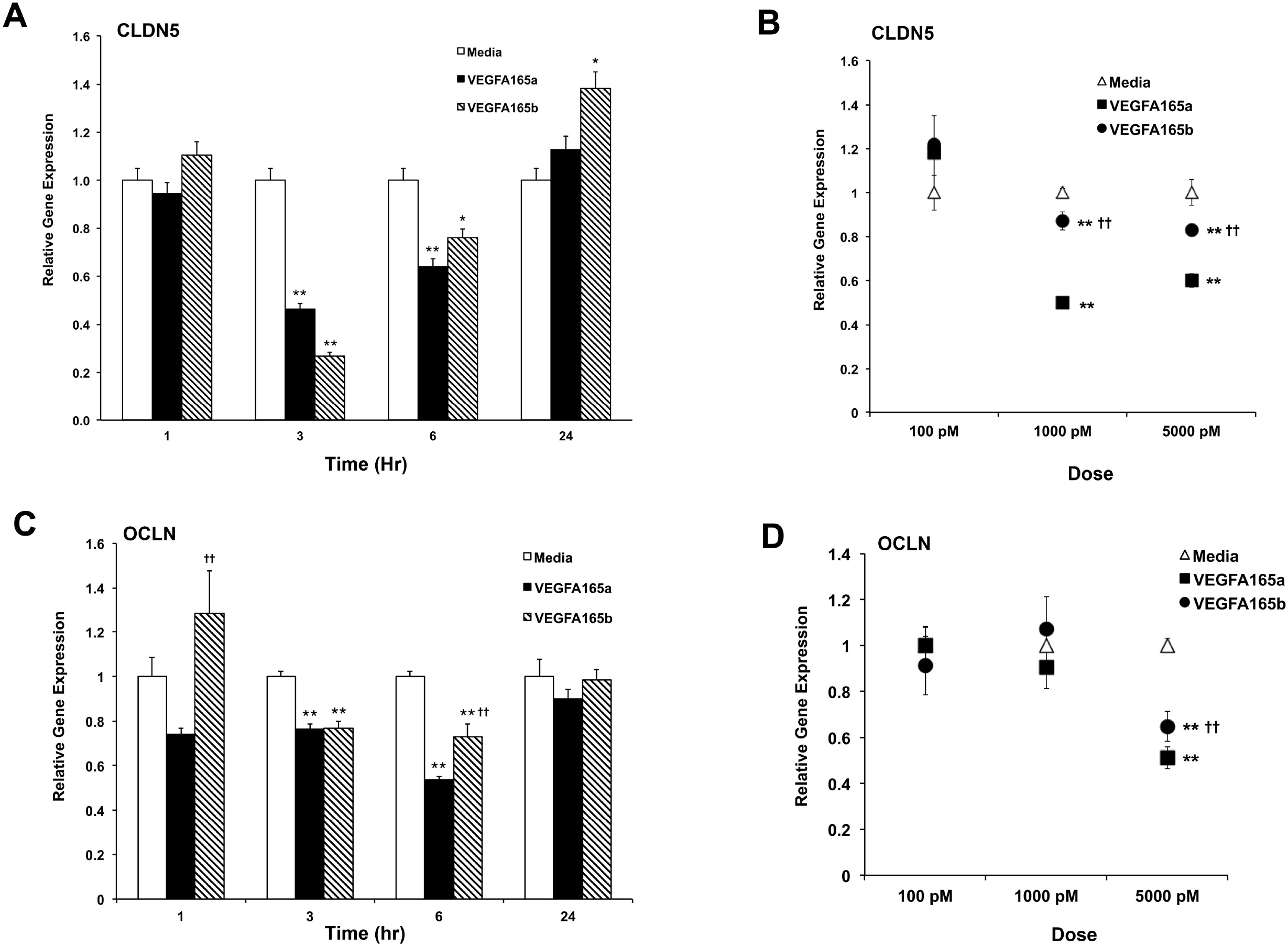
VEGFA_165_a has a stronger effect on tight-junction gene expression in primary HRMECs than VEGFA_165_b. Relative gene expression measured by qPCR. **A)** *Claudin-5* gene expression at 1, 3, 6 and 24 hours after VEGFA_165_ treatment. Confluent HRMECs were treated with a saturating high-dose of VEGFA_165_a or VEGFA_165_b (100 ng/mL, 5300 pM). **B)** *Claudin-5* gene expression comparing VEGFA_165_a and VEGFA_165_b at a low, intermediate and high doses measured after 6-hours. **C)** *Occludin* gene expression at 1, 3, 6 and 24 hours after VEGFA_165_ treatment. **D)** *Occludin* gene expression comparing VEGFA_165_a and VEGFA_165_b at a low, intermediate and high dose measured after 6-hours. (Triplicate assays, error bars show standard deviation. Anova: compared to media control *p<0.05, **p<0.01; compared to VEGFA_165_a ^††^ p<0.01)

## DISCUSSION

While the role of VEGFA in vascular development, tumorgenesis and hypoxia has been studied for almost three decades, less is known about the functional differences between VEGFA isoforms. More recently, isoform-specific analysis reported that most VEGFA_165_ in normal vitreous was VEGFA_165_b, with the ratio changing to mostly VEGFA_165_a in eyes with active diabetic retinopathy and ROP [33,34]. The relative expression of VEGFA_165_a and VEGFA_165_b can be regulated by alternative splicing of the VEGFA_165_ pre-mRNA, which is regulated by phosphorylation of the ASF/SF2 splice-factor by SR-protein kinase (SRPK1/2) [39]. This raised the possibility that modulation of this isoform ratio might be exploited for therapeutic purposes. Modulation of the splicing between the VEGFA_165_a and VEGFA_165_b mRNAs was demonstrated using SRPK inhibitors in human retinal pigment epithelial (RPE) cells, and topical infusion of these inhibitors reduced neovasularization in a mouse laser-induced CNV model [40]. Another group used intravitreal injection of a different SRPK inhibitor to inhibit neovascularization in the mouse CNV model [41]. An antibody specific for the C-terminus of VEGFA_165_a, which does not bind VEGFA_165_b, can also inhibit the proangiogenic effects of VEGFA_165_a [42].

Specific blockade of the VEGFA_165_a protein, or supressing VEGFA_165_a expression in favor of VEGFA_165_b, could be a therapeutic strategy if there are significant differences in the ability of these isoforms to activate intracellular signalling in the retinal endothelium. The preference for examining signaling differences using primary HRMECs is supported by studies that revealed differences between endothelial cells derived from different organs. For example, viral delivery of a murine recombinant-VEGF_164_b attenuated inflammatory-response damage in a murine model of ulcerative colitis, yet it exacerbated blood-brain-barrier damage in a model of focal cerebral ischemia [43-45]. Extensive gene expression profiling has also demonstrated that HUVECs have a greater expression of embryonic genes, and other differences, compared to endothelial cells from the choroid and neural retina. Significant differences also exist between human retinal endothelial cells and human choroidal endothelial cells in the expression of genes that are related to endothelial function and neovascularization [46].

We began with immunoblotting to examine the activation of MAPK and AKT in primary HRMECs. While VEGFA_165_b does not bind the Neuropilin-1 co-receptor, it does bind to VEGFR2 with similar or less affinity as VEGFA_165_a depending on the cell model and method used [31,32]. Irregardless of relative binding affinity, VEGFA_165_b is reported to bind and activate VEGFR2 based on immunoblotting studies [31,47,48]. Using immunoblotting, we found that both MAPK and AKT were activated to a similar extent by VEGFAM_165_a or VEGFA_165_b, using a dose of 100 ng/ml (5,300 pM) in primary HRMECs. Previous single-dose immunoblotting studies in different cell-types have reported that VEGFA_165_b activated VEGFR2 and MAPK less than VEGFA_165_a, and that the maximum activation of MAPK by VEGFA_165_b occurred later than VEGFA_165_a in porcine aortic endothelial cells (PAECs) and bovine aortic endothelial cells using a dose of 2,600 pM [31,47]. Another study of human pulmonary microvascular endothelial cells (HPMECs) found that the a-isoform was a stronger activator of VEGFR2 and MAPK (ERK1/2) than the b-isoform at a dose of 20 ng/ml (1000 pM) [48].

We found that both MAPK and AKT were activated less by VEGFA_165_b at the lower dose (20 ng/ml, 1000 pM) (data not shown). However, it was noted that while activation was detected in repeated experiments, the relative manitude of activation was variable using immunoblotting.

Immunublotting does involvenumerous processing steps and a subtantial number of cells are required for a single sample, therefore we decided that *in situ* assays would be supperior for generating full dose-response curve data.

To determine the dose-response for activation of MAPK and AKT, *in situ* ICW assays were developed that permited the use of multiple replicate doses with a limited supply of primary cells in a 96 well plate format. The resulting dose-response curves confirmed that there were substantial differences in the activation of both MAPK and AKT between VEGFA_165_a and VEGFA_165_b in primary HRMECs. In the case of MAPK, the ED_50_ of VEGFA_165_a 900 pM less than the ED_50_ for VEGFA_165_b. VEGFA_165_a caused near maximal activation of MAPK at a concentration of 250 pM, whileVEGFA_165_b had little effect at 250 pM. Higher doses of VEGFA_165_b could activate MAPK to a similar maximum level, and the timing for maximum activation of MAPK was similar for both isoforms in HRMECs, occuring by 10 minutes and decreasing by 30 minutes. This relative timing was different from that reported for HUVECs where the maximum concentration of active MAPK occured 30 minutes after treatment with VEGFA165b [31].

For AKT, the difference in ED_50_ between VEGFA_165_a and VEGFA_165_b was not as large as seen for MAPK, just two-fold greater for VEGFA_165_b compared to VEGFA_165_a. However, the activation of AKT by VEGFA_165_b was subtanially less than from VEGFA_165_a over the entire dose-response range. In contrast to MAPK activation, the time to maximum activation of AKT was earlier with VEGFA_165_b (15 minutes) compared to the VEGFA_165_a (30 minutes). While the differences in ED_50_ values were not as large as seen for MAPK activation, the maximum activation of AKT obtained with VEGFA_165_b was only 50% of the maximum activation generated by VEGFA_165_a. This was another difference from that reported in a non-retinal endothelial celltype, human pulmonary endothelial cells (HPECs), whereAKT activation is the same for VEGFA_165_a and VEGFA_165_b at a dose of 20 ng/ml (1000pM) in HPECs [48].

The differences noted above in the activation of MAPK and AKT between HRMECs and non-retinal endothelial cells suggest that VEGF-mediated intracellular signaling patterns are not completely universal between endothelial cells from different organs. An evaluation of primary endothelial cells from the human retina should be utilized whenever possible to study the retinal context. Altogether, our dose-response curves for the activation of VEGFR2, MAPK, and AKT in primary HRMECs were consistent with the model proposed by Whitaker et al. (2001), wherein VEGFA_165_a activates VEGFR2 more intensely when it can bind the co-receptor Neuropilin-1, to form a larger activation complex [49].

Based on the primary HRMEC dose-response data for MAPK activation, VEGFA_165_a was more potent for activating intracellular signalling than VEGFA_165_b at lower doses, which fall into a range of total VEGFA_165_a concentrations previously reported in human vitreous. Vitreous concentrations for VEGFA have been measured by various techniques in several disease conditions. Most elevated VEGFA is the VEGFA_165_ isoform, which diffuses slowly, and the vitreous has protease activity, therefore it is possible that the concentrations measured in the vitreous are somewhat lower than in the neural retina itself. With that stated, vitreal concentrations of 25 pM to over 400 pM have been reported for proliferative diabetic retinopathy from several studies [1,50–54]. Other studies have reported 10 pM in diabetic macular edema [55], 100 to 450 pM in retinopathy of prematurity [34,56] and an average of 430 pM, with as high as 580 pM, in central retinal vein occlusion [57]. It is possible that elevations of retinal VEGFA_165_a concentration to several-hundred pico-Molar could increase the activation of endothelial intracellular signalling above its normal baseline. In contrast, the much lower activation potential of VEGFA_165_b in this concentration range would leave MAPK activation close to normal if most of the VEGFA_165_ existed as the b-isoform.

We also found that VEGFA_165_b could affect the expression of the same target genes that are also regulated by VEGFA_165_a in primary HRMECs. These included genes involved in leukocyte-endothelial cell adhesion and tight-junction formation. Even though the 5,000 pM dose of VEGFA_165_b could activate MAPK and AKT more strongly than the intermediate 1,000 pM dose, it is unlikely that the 5,000 pM concentration would be experienced *in vivo*. For that reason we suggest that the effects on gene expression at 1,000 pM were most informative. We found that the *CLDN5* gene was particulary succeptible to repression by both isoforms of VEGFA_165_ compared to *OCLN* in this cell type. VEGFA-mediated activation of the VEGFR2 receptor is known to disrupt adherin-junctions and causes ß-Catenin mediated repression of *Claudin-5* gene expression in endothelial cells [58]. For all three leukocyte docking protein genes (*ICAM1, SELE, VCAM1*), VEGFA_165_a increased their expression at the intermediate dose of 1000 pM where VEGFA_165_b had less effect. We concluded that there were significant dose-response differences in the regulation of VEGFA-target genes, with VEGFA_165_a proving to be the stronger affector of gene expression overall.

### Summary

Our results indicated that differences in the last 6 amino acids between VEGFA_165_a and VEGFA_165_b have a significant affect on the relative activation of MAPK and AKT within primary HRMECs. VEGFA_165_a activated both pathways at lower concentrations where VEGF_165_b had little effect. The greater potential of VEGFA_165_a to affect HRMECs also extended to changes in the expression of genes required for leukocyte-docking and tight-junction structure. Large dose-response differences in activation of these pathways exist in ranges of elevated VEGFA_165_a concentrations that have been reported in the vitreous of diseased eyes. Key aspects of MAPK and AKT activation, such as timing and maximal activation, were different in HRMECs compared to those reported for some other non-retinal endothelial cell-types. These results would support the concept that specific blockade of VEGFA_165_a, or modulating the VEGFA_165_b / VEGFA_165_a ratio, could be useful thereapeutic strategies for retinal diseases involving elevated VEGFA concentration.

## Supporting information

Supplemental Observations Rat Eye

## Acknowledgements

The authors would like to thank Oakland University students Regan Miller and Anju Thomas for assistance with some cell culture and some testing of gene expression probesets used for these studies. Research supported by National Eye Institute / National Institutes of Health (USA) grant NIH EY025089 (KPM).

