## Supplemental Observations Rat Eye for "Differences in Activation of Intracellular Signalling in Primary Human Retinal Endothelial Cells Between Isoforms of VEGFA-165"

While testing a VEGFA**_165_**a intraocular-injection rat model, which causes primary retinal vein dilation and tortuosity [1], the unexpected observation was made that VEGFA**_165_**b had the same effect as VEGFA**_165_**a. While VEGFA intra-vitreal injection is used to reduce vascular permeability, the precise control of concentration is difficult in the vitreous and saturating doses are generally used for effect. While not the main subject of the manuscript, regarding activation of intracellular signaling by VEGFA**_165_**a and VEGFA**_165_**b, the following observations in vivo inspired the pursuit of investigations with primary human retinal endothelial cells.

Examples of the observations and the imaging methods for detecting changes in retinal vein diameter and neural retinal edema are provided here for readers who interested in these methods for their own research. The procedures include guidance for rat anesthesia, pupil dilation, fluorescein angiography, Evans Blue angiography, SD-OCT, and measurement of retinal thickness at multiple locations in a single retina using DIVER software with OCT data.

**Ocular Imaging Methodology**

*Animal care and use:* All experiments performed in this study were carried out with the approval of Oakland University’s Animal Care and Use Committee and conformed to the ARVO Statement for the Use of Animals in Ophthalmic and Vision Research. Rats were housed at Oakland University in a facility approved by the Association for Assessment and Accreditation of Laboratory Animal Care International. Adult Long Evans rats (155-190 grams body weight, female) were obtained from Charles River Laboratories (Wilmington, MA).

*Anesthesia for injections or retinal imaging:* Pupils were dilated with tropicamide and phenylephrine drops prior to anesthesia. Rats were anesthetized with an intra-peritoneal injection of a Ketamine HCL (50 mg/kg) and Xylazine (7 mg/kg).

*Fluorescein Angiography:* After sedation and pupil dilation, rats were injected (intra peritoneal) with 50 microliters of 10% sodium fluorescein in PBS. Retinas were imaged using a MICRON 3 camera system using the fluorescein filter set (blue illumination, green emission). Corneal surfaces were covered using GenTeal lubricant eye gel (Novartis, CVS Pharmacy) to prevent corneal dehydration and to provide optical transmission with the camera lens. Images were captured using a camera system gain of 12 with 20 frame averaging to allow for low energy illumination. OD eyes were imaged first, then the same illumination setting was used to switch to imaging the contralateral OS eye.

*VEGFA_165_a and VEGFA_165_b intra-vitreal injection:* All animals were pre-imaged, prior to injections of VEGFA, using fluorescein angiography and SD-OCT imaging to record the status of their normal retinal vasculature. Both imaging modes were completed in a single session of anesthesia. This was typically performed about one week before intraocular injections with VEGFA isoforms. Rats were anesthetized and their pupils dilated prior to intraocular injections. Recombinant carrier-free human VEGFA_165_a, or VEGFA_165_b, was reconstituted with phosphate buffered saline solution (PBS, no calcium, no magnesium) to the appropriate concentration for intra-vitreal injections of a 5μl volume (dose 300 ng/eye) for saturation. Injections were done with a NanoFil syringe fitted with 35 gauge and beveled NanoFil needles (WPI Inc., Sarasota, Florida). The contralateral eye served as a control and received the same injection procedure-using vehicle alone (PBS). Depending on the experimental goal, imaging of the retinal vasculature was generally repeated at 24, 48, or 72 hours post injection and retinas collected for biochemical analysis.

*Analysis of rat retinal edema and primary retinal vein diameter in vivo:* Spectral Domain Optical Coherence Tomography (SD-OCT) was utilized to measure changes in retina thickness *in vivo*, induced by intra-vitreal injection of VEGFA isoforms. Following the convention, OD eyes were injected with VEGFA**_165_**a or VEGFA**_165_**b, while OS eyes received an injection of vehicle (PBS). To maintain corneal transparency for imaging, Artificial Tears lubricant eye drops (CVS Pharmacy) solution were applied generously to the both corneal surfaces. Anesthetized rats were secured in a three-axis positioning support cradle (Bioptigen, Durham NC). SD-OCT scans were taken using an Envisu R2200 model SD-OCT system (Bioptigen) equipped with a lens for the rat eye axial length. A rectangular scan pattern of rat sizes 2.6 mm x 2.6 mm was used (1000 A-scans by 100 B-scans). For measuring retinal vein thickness, calipers were positioned and locations recorded for subsequent re-measurements of the same primary veins locations after injection of VEGFA isoforms. Three to four veins were monitored this way per retina to determine an average percent change in diameter between the pre- and post-injection time points.

To measure edema related thickening of the retina, the thickness of the neural retinal vascular compartment was measured between the Inner Retinal-Nerve-Fiber-Layer (Inner RNFL) and the Outer Outer-Nuclear-Layer (Outer ONL) using Bioptigen’s InVivoVue Diver 2.4.10 software [2]. An example of software use for the rat retina is illustrated in Supplemental Figure-1. A fixed 5x5 grid was first centered on the optic disc and the boundaries of all retinal layers were marked at all grid locations. The central grid position was not used for measurements as it was used for centering on the optic disc center. The remaining 24 grid positions were then used to calculate an average thickness for each retina, pre- and post-injection. Thickness was also compared by averaging the 8 most central grid locations and the 16 most peripheral grid locations.

*Evans Blue Angiography:* The same animals utilized for layer analysis with DIVER software were also imaged by Evans Blue angiography to detect leakage into the vitreal chamber in treated eyes (OD) versus vehicle (PBS) injected control eyes (OS). Fluorescein angiography images were obtained several days prior to intravitreal injections to confirm a normal retinal vasculature. 24-hours after VEGFA**_165_** isoform injections all eyes were imaged again using Evans Blue angiography. Evans Blue solution (30mg/ml) at a dose of 30 mg/kg was injected intravenously (tail vein) two hours prior to imaging for Evans Blue Angiography. After sedation and pupil dilation, retinas were imaged using a MICRON-3 camera system (Phoenix technology group, Pleasanton, CA) using the Evans Blue filter set (green illumination, red emission). Corneal surfaces were covered using GenTeal lubricant eye gel (Novartis, CVS Pharmacy) to prevent corneal dehydration and to provide optical transmission with the camera lens.

**Supplement Figure-1. Method for measurement of retinal layer thickness in the rat eye.** After loading of a 3-dimensional SD-OCT data set, a 5x5 grid was centered on the optic disc using the grid panel (left). The series of b-scans located at each selected grid position (red circle) were viewed and marked for retinal layer locations as shown in the center panel, top. The Inner Retinal-Nerve-Fiber-Layer (Inner RNFL), Outer Outer-Plexiform-Layer (Outer OPL) and Outer Outer-Nuclear-Layer (Outer ONL) are labeled here for orientation. Thickness of the neural retina vascular compartment was measured as the difference between the Inner RNFL and the Outer ONL, which mark the locations of the inner and outer membranes. This thickness was determined for each retina as an average from all grid locations, as well as averages from the just the most central grid positions (8, in the white box) and the peripheral grid positions (16).

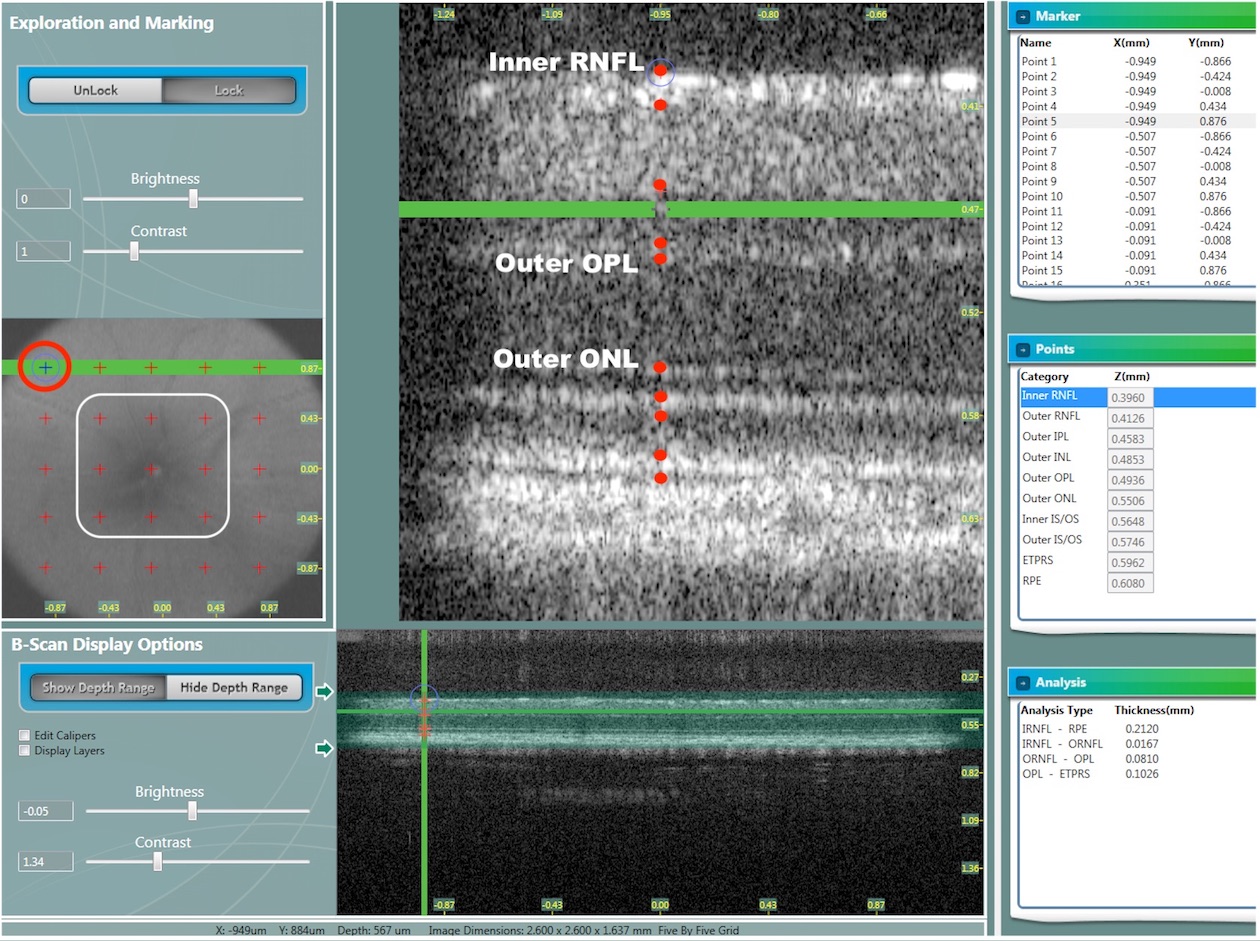

**Supplement Figure-2. Examples primary retinal vein dilation observed 24 hours after intra-vitreal injection of VEGFA_165_a or VEGFA_165_b.**

Both VEGFA**_165_**a and VEGFA**_165_**b caused visible dilation of the primary retinal veins. Vehicle (PBS) injected control eyes did not show vein dilation. Fluorescein angiography images of eyes were obtained before injection and 24 hours after injection with isoforms of A) VEGFA**_165_**a. Right (OD) eyes were injected with 250 ng VEGFA**_165_**a or and contralateral (OS) eyes were injected with vehicle (PBS) as controls. B) VEGFA**_165_**b. Right (OD) eyes were injected with 250 ng VEGFA**_165_**b or and contralateral (OS) eyes were injected with vehicle (PBS) as controls. Primary veins marked (v).

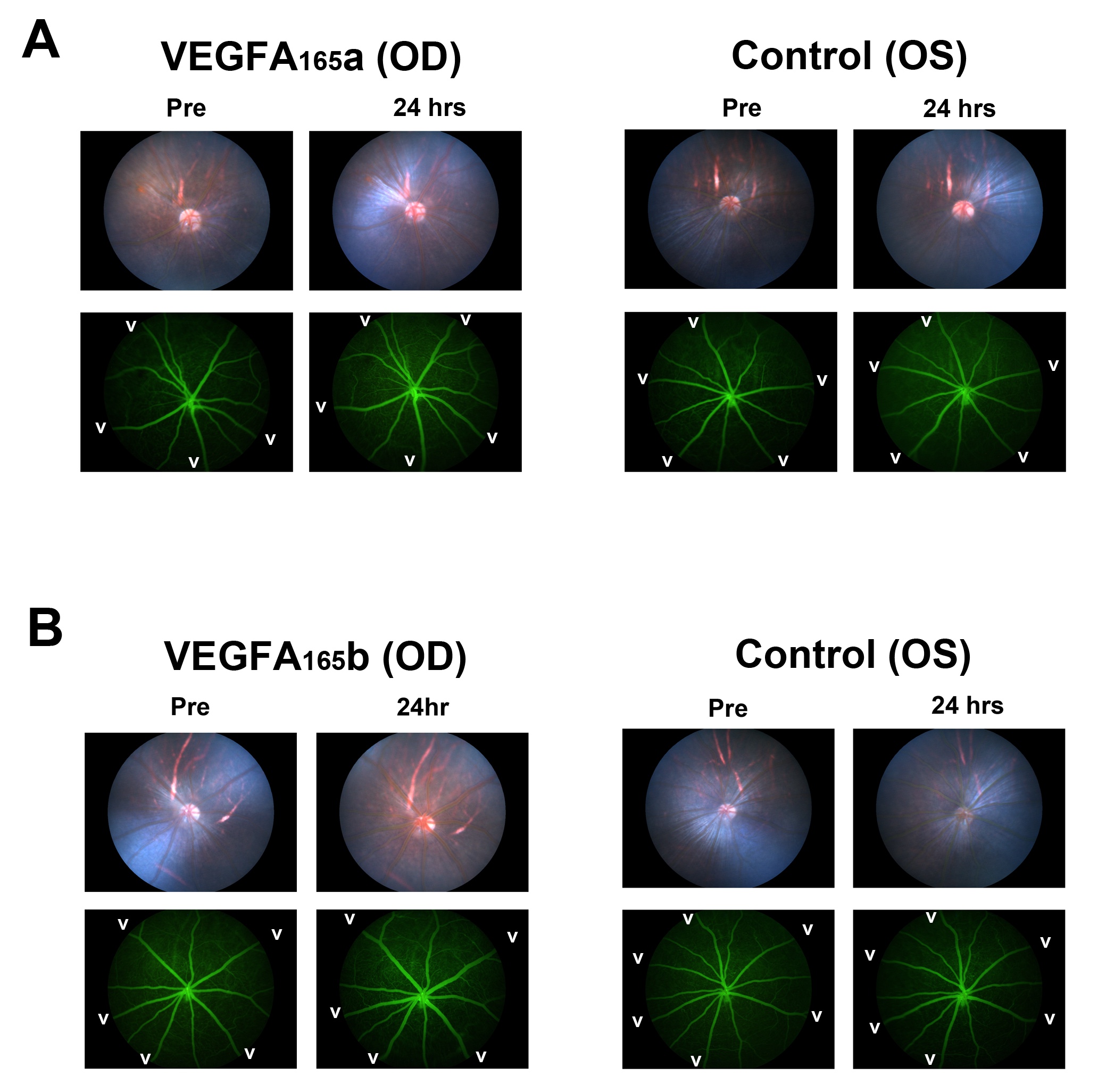

**Supplemental Figure-3. Observation of primary vein dilation at 24 hours post-injection and recovery to pre-injection diameter by 72 hours following intra-vitreal injection of VEGFA_165_b.**

Primary vein diameter followed by fluorescein angiography and SD-OCT images in the same rat retina imaged before and after intra-vitreal injection of VEGFA165b (250 ng).

**A)** Fluorescein angiography images. All five primary retinal veins displayed a visible dilation at 24 hours post-injection. Recovery to pre-injection diameter was essentially complete by 72 hours post-injection. **B)** SD-OCT *en face* views of primary vessel shadows. The same vein as in panel-A are marked with a black (v). The green horizontal lines and red-arrows indicate the location of the zoomed OCT cross-sectional images shown in panel-C. **C)** Zoomed view of OCT cross-sectional images with digital-calipers to indicate the diameter of the vein at the location (v) in panel-B.

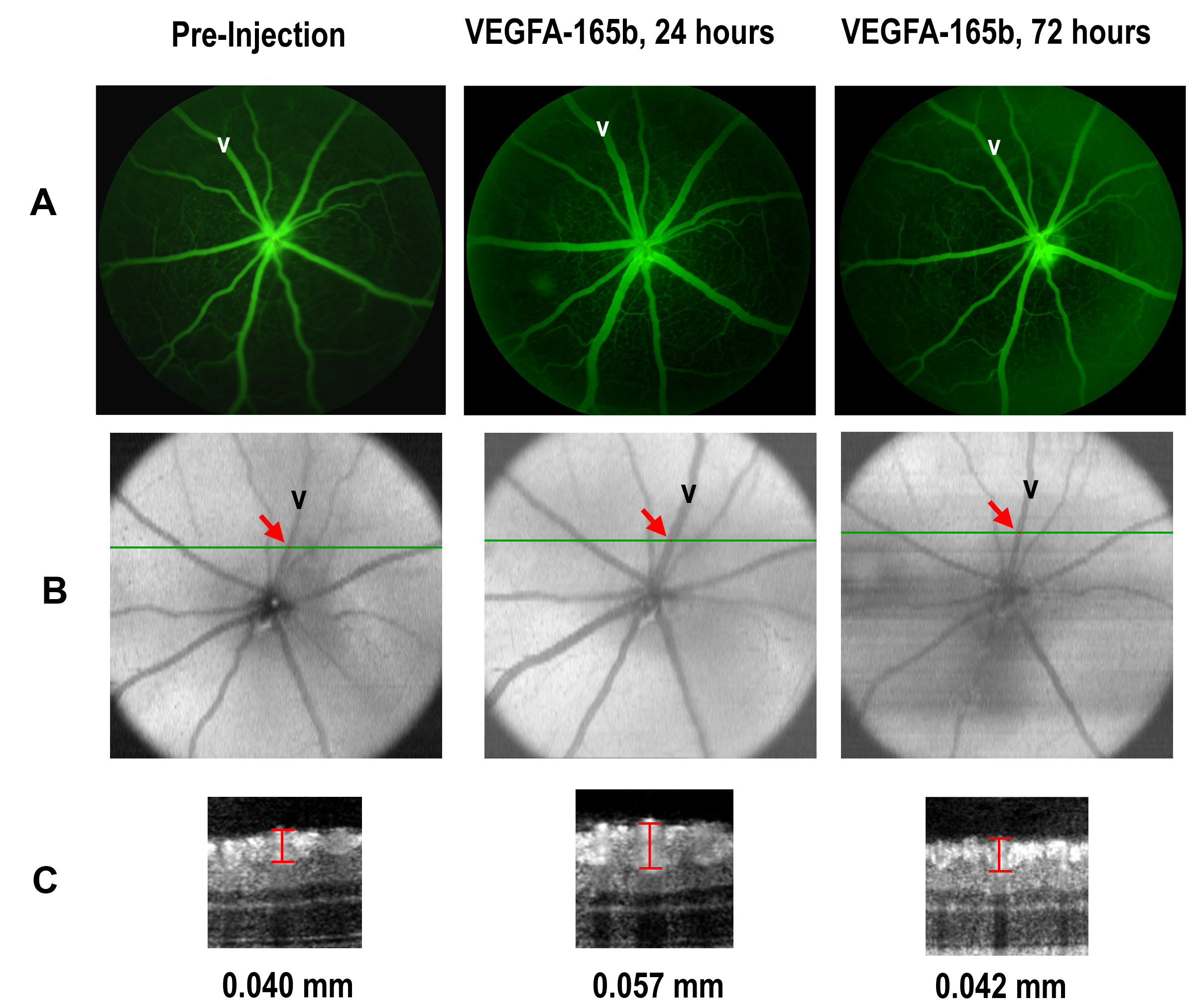

**Supplemental Figure-4. Control observation that the VEGF-trapping drug Aflibercept could block the primary retinal vein dilation caused by VEGFA_165_b.**

Fluorescein angiography, showing the dilation of primary retinal veins 24 hours post injection with VEGFA**_165_b** (OD eye, white arrows). Significant recovery from the transient dilation is seen at 48 hours post-injection. Co-injection of Aflibercept with VEGFA**_165_b**, in the OS eye, blocked the dilation effect at 24 hours.

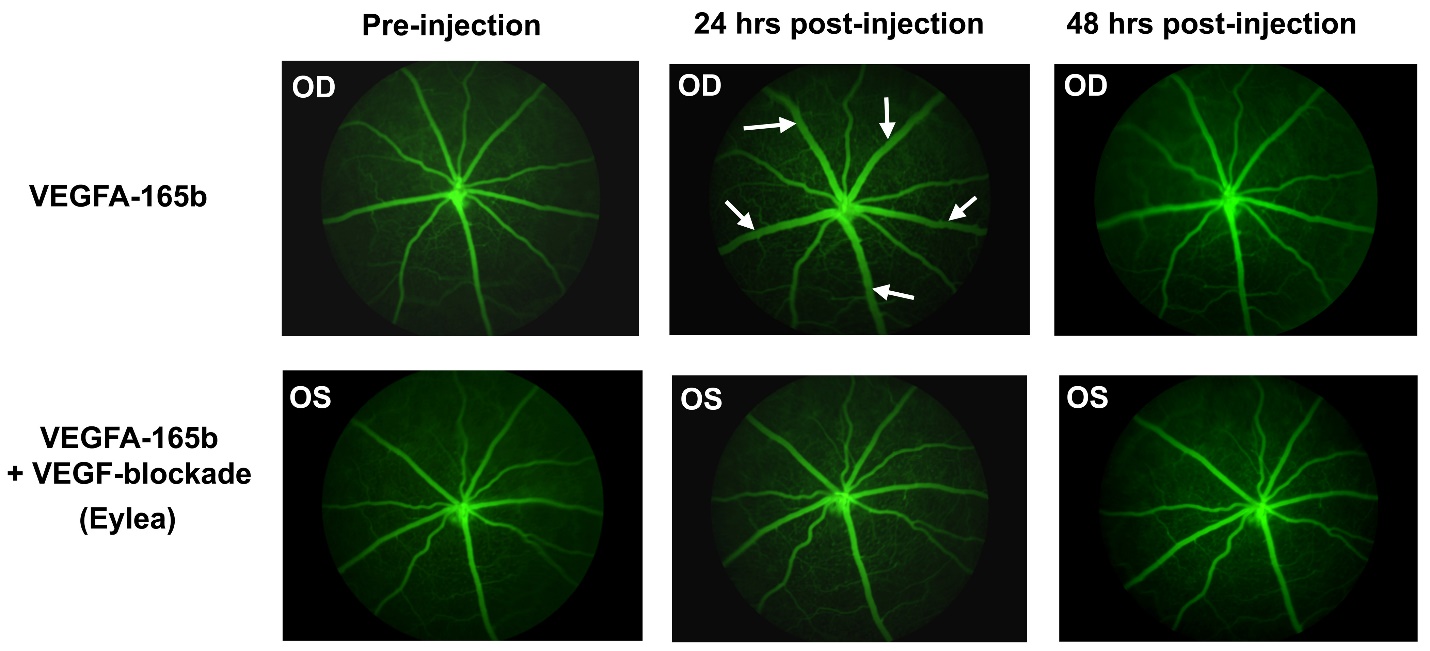

**Supplemental Figure-5. Changes in the diameter of primary retinal veins by VEGFA_165_a and VEGFA_165_b *in vivo*.**

Rats received intra-vitreal injections in one eye with either VEGFA**_165_**a or VEGFA**_165_**b (300ng). The contralateral eye received an intra-vitreal injection with PBS (vehicle) only. Retinas were imaged by SD-OCT prior to injection and again at 24 hours post-injection. The average percent diameter change from pre- to post-injection was calculated using three primary vein diameter measurements per retina. (*p<0.03 vs vehicle, **p<0.005 vs vehicle. N=2 rats used for each isoform)

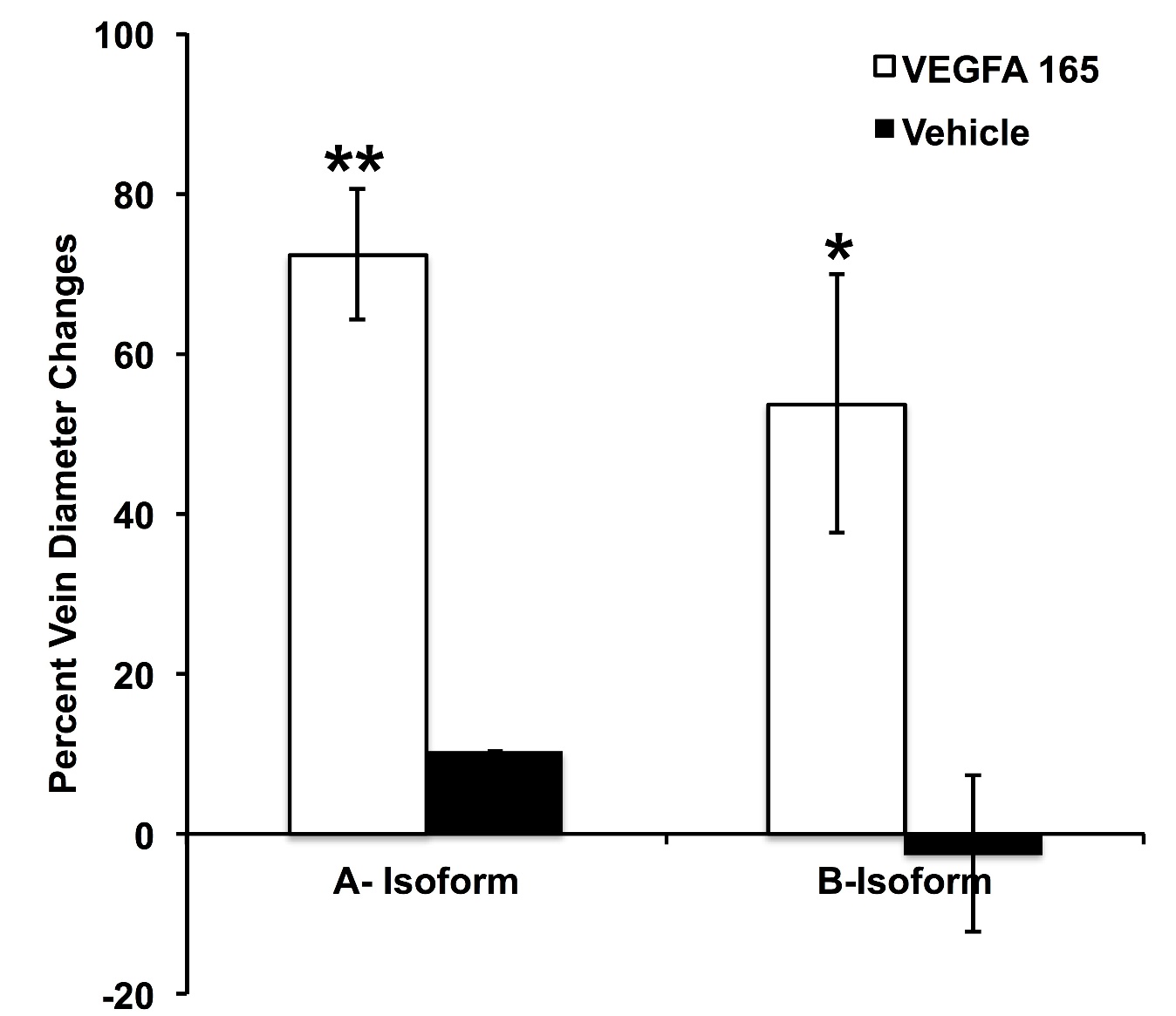

**Supplemental Figure-6. Edema of the neural retinal vascular compartment induced by by VEGFA_165_a and VEGFA_165_b.**

Thickness of neural retinal vascular compartment, bounded by the Inner-Limiting Membrane (ILM) and Outer-Limiting Membrane (OLM), was measured from SD-OCT data using DIVER software. Thickness was averaged from 24 measurements per retina, centered on the optic disc and including the central to mid retina. (Refer to Supplemental Figure-1) Central thickness was averaged from 8/24 locations adjacent to the optic disc. Mid thickness was averaged from 16/24 locations more peripheral to the optic disc. **A)** VEGFA**_165_**a caused a relative increase in thickness in OD eyes compared to PBS-injected OS eyes averaged over the total area, central area, or mid area. **B)** VEGFA**_165_**b also caused a relative increase in thickness compared to PBS-injected contralateral eyes for the total area, central area and mid area. (t-test, N=2 rats per group, * P<0.05, ** P<0.01)

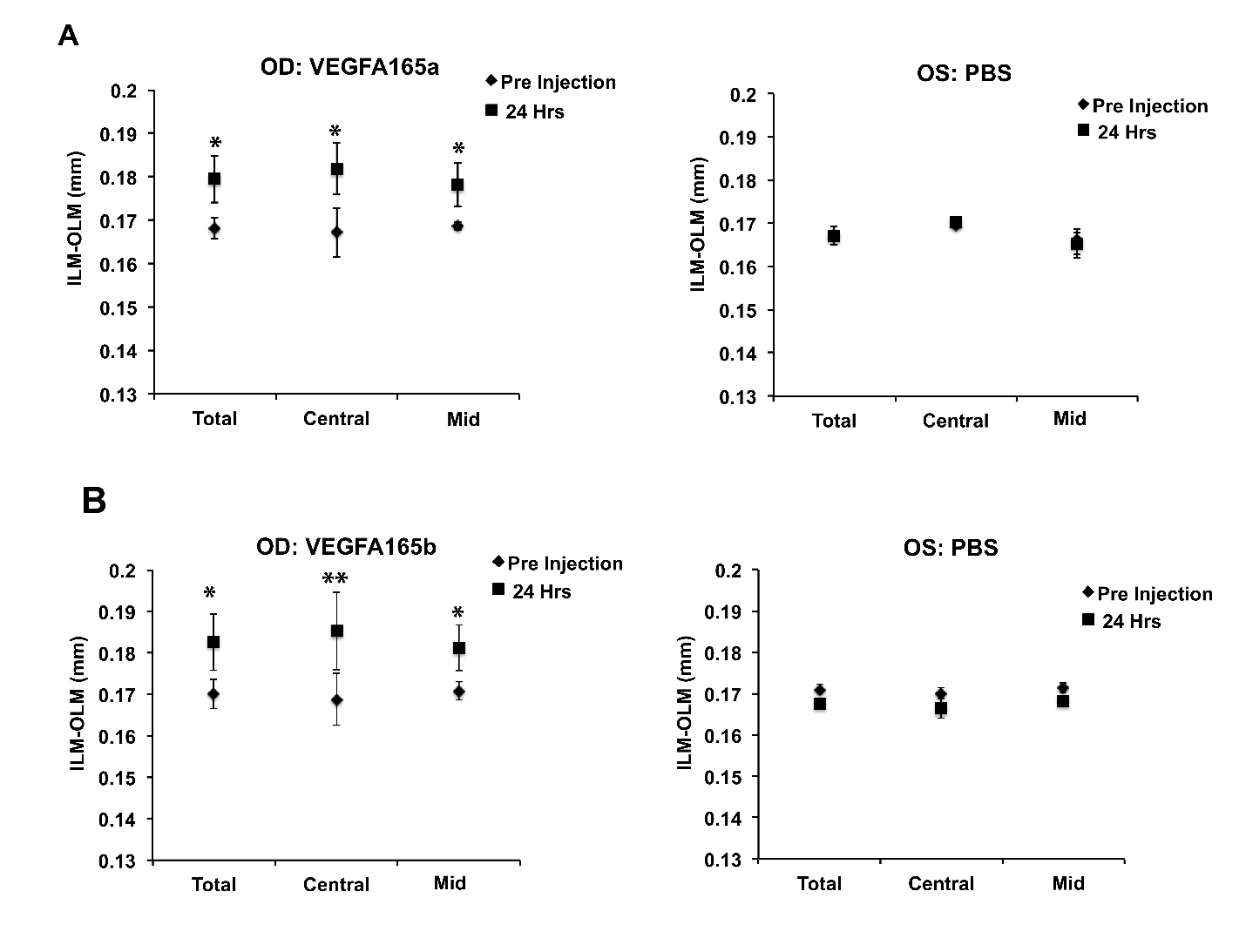

**Supplemental Figure-7. Increased leakage of intravenous Evans Blue dye into the vitreous humor of eyes injected with VEGFA_165_a or VEGFA_165_b.**

Retinas from the same rats used for measurements of thickness changes (Supplemental Figure-6) were imaged by fluorescein angiography before injection of VEGFA**_165_** isoforms. 22-hours post-VEGFA injections (intra-vitreal), animals received an intravenous injection of Evans Blue dye. At 24 hours retinas were imaged by Evans Blue angiography. Treatment of right eyes (OD) with **A**) VEGFA**_165_**a or **B**) VEGFA**_165_**b increased leakage of Evans Blue into the vitreous compared to PBS-injected left eyes (OS).

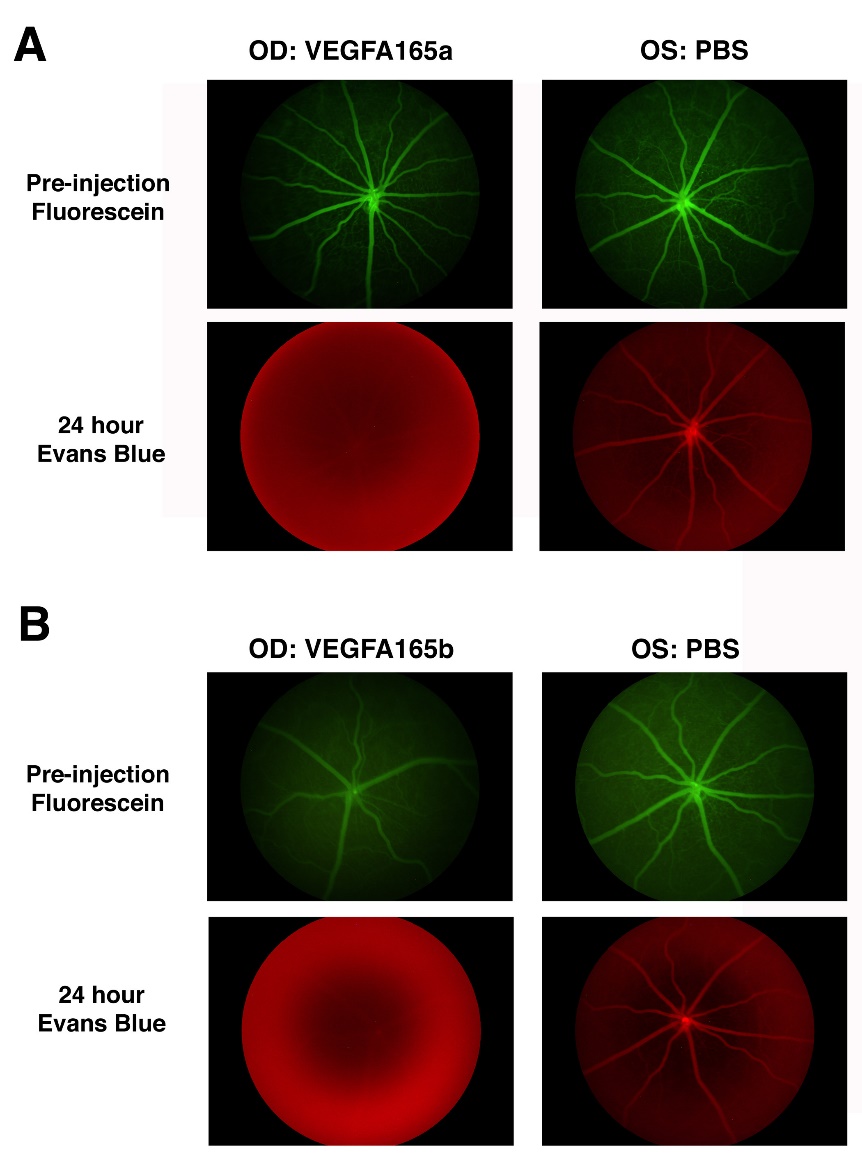
